# Evolutionary dynamics of SARS-CoV-2 nucleocapsid protein (N protein) and its consequences

**DOI:** 10.1101/2020.08.05.237339

**Authors:** M. Shaminur Rahman, M. Rafiul Islam, A. S. M. Rubayet Ul Alam, Israt Islam, M. Nazmul Hoque, Salma Akter, Md. Mizanur Rahaman, Munawar Sultana, M. Anwar Hossain

## Abstract

The emerging novel coronavirus SARS-CoV-2 has created a global confusing pandemic health crisis that warrants an accurate and detailed characterization of the rapidly evolving viral genome for understanding its epidemiology, pathogenesis and containment. We explored 61,485 sequences of the Nucleocapsid (N) protein, a potent diagnostic and prophylactic target, for identifying the mutations to review their roles in RT-PCR based diagnosis and observe consequent impacts. Compared to the Wuhan reference strain, a total of 1034 unique nucleotide mutations were identified in the mutant strains (49.15%, n=30,221) globally. Of these mutations, 367 occupy primer binding sites including 3’-end mismatch to primer-pair of 11 well characterized primer sets. Noteworthy, CDC (USA) recommended N2 primer set contained lower mismatch than the other primer sets. Moreover, 684 amino acid (aa) substitutions located across 317 (75.66% of total aa) unique positions including 82, 21, and 83 of those in RNA binding N-terminal domain (NTD), SR-rich region, and C-terminal dimerization domain (CTD), respectively. Moreover, 11 in-frame deletions were revealed, mostly (n =10) within the highly flexible linker region, and the rest within the NTD region. Furthermore, we predicted the possible consequences of high-frequency mutations (≥ 20) and deletions on the tertiary structure of the N protein. Remarkably, we observed that high frequency (67.94% of mutated sequences) coevolving mutations (R203K and G204R) destabilized and decreased overall structural flexibility. Despite being proposed as the alternate target to spike protein for vaccine and therapeutics, ongoing nonsynonymous evolution of the N protein may challenge the endeavors, thus need further immunoinformatics analyses. Therefore, continuous monitoring is required for tracing the ongoing evolution of the SARS-CoV-2 N protein in prophylactic and diagnostic interventions.

## 1. Introduction

In December 2019, viral pneumonia cases of unknown cause emerged in Wuhan City, China (Liu et al., 2020; Mo et al., 2020). Later, the scientists confirmed the etiology as severe acute respiratory syndrome coronavirus 2 (SARS-CoV-2); and named the disease as COVID-19 (Mo et al., 2020; Shi et al., 2020). The entire world has witnessed the rapid spread of this virus causing thousands of mortalities each day and creating an unprecedented level of public health emergency (Gussow et al., 2020; Islam et al., 2020a). Among four structural proteins of this virus, the N protein is considered as the most important multifunctional RNA binding protein involved in several aspects of virus life cycle such as ribonucleoprotein helical structure formation during packaging of the genome, RNA synthesis regulation during replication, transcription and modulating cellular metabolism of the host (Kang et al., 2020; Liang et al., 2020). In addition, the N proteins of coronaviruses are highly expressed in the host cell, and involved in regulating host-pathogen interactions during infection (Liang et al., 2020). Moreover, this protein has a high immunogenic property enabling it to elicit protective immune response against SARS-CoV-2 (Kang et al., 2020; Liang et al., 2020). However, the molecular mechanism involving N protein in pathogenesis is yet to be clearly understood.

The N protein is the only structural protein present in the nucleocapsid of the SARS-CoV-2 genome which is 419 amino acid (aa) long, and composed of three distinct domains: N-terminal domain (NTD)/ RNA-binding domain (46-176 aa), a serine/arginine-rich (SR-rich) (184-204 aa) linker region (LKR) (182-247 aa), and a C-terminal domain (CTD) (247-364 aa) (Kang et al., 2020; Liang et al., 2020; Zeng et al., 2020). The NTD binds to the 3′-end of the viral RNA, and is highly divergent. The charged serine and arginine-rich LKR region can interact directly with RNA, and play a part in cell signaling (Hurst et al., 2009; Liang et al., 2020). The N-NTD, N-CTD, and SR-rich domain have been found to be associated with genomic RNA of the virus through electrostatic interactions by the positively charged aa residues and modulate unwinding after entering into cell by phosphorylation to specific aa. In addition, several critical residues of N protein have been identified to be involved in genomic RNA binding, and virus infectivity modulation (Cascarina and Ross, 2020; Zeng et al., 2020). Interestingly, Lin et al. (2020) has recently investigated the fate of N protein expression in human induced pluripotent stem cells (iPSC) and observed that it abolished pluripotency and reduced the proliferation rate whereas long-term expression the protein drives iPSC to fibroblast (Lin et al., 2020). Since this novel virus shows a distinct electrostatic surfaces and topological variations in the β-hairpin of NTD even though resembles similar structures with closely related coronaviruses (Zhou et al., 2020), a detailed analyses on structural dynamics due to significant mutations can likely answer whether they have any specific adaptive functions.

Mutations have been found to occur in all regions of the virus, including the open reading frames (ORFs), the S protein, and the N protein (Islam et al., 2020a; Ozono et al., 2020). Several types of substitutions in different types of viral proteins have already been reported within a short period of time that might be involved in modulating viral transmissibility, replication efficiency as well as virulence properties of the virus in different parts of the world (Islam et al., 2020a; Jia et al., 2020; Pachetti et al., 2020). The COVID-19 symptoms cannot be used for accurate diagnosis as patients express nonspecific symptoms, and many of these symptoms could be associated with other respiratory illnesses. Recently molecular techniques for nucleic acid detection have been used for diagnosing and screening COVID-19 (Udugama et al., 2020). A number of different primer-probe sets targeting ORF1ab, S, and N genes have been used in SARS-CoV-2 detection assays, hence COVID-19 detection testing kits have been well developed, and are now available worldwide (Nalla et al., 2020; Park et al., 2020). However, the test results can be affected by the nucleotide variations present on the primer binding sites of viral RNA sequences. Mutations in the primer binding sites of the N gene affects the assay performance since it largely depends on the sequence matching between primer-probe and SARS-CoV-2 sequences, mismatches of which can cause the inconsistency or the high false negative rate (FNR) in the test results (Nalla et al., 2020; Rainer et al., 2004, 2020). Moreover, ongoing mutational evolution of the virus warrants continuous monitoring and updating of primer-probe sets targeting the indigenous genome sequences for specific and accurate detection.

In this study, we have pursued an investigation into the synonymous and nonsynonymous mutations across 61,485 full-length N protein gene trimmed from the SARS-CoV-2 genome sequences. Our targets are to observe any nucleotide change that can cause the mismatching of recommended RT-PCR primer-probe and identify the aa substitutions in the protein that might cause alteration in the structural configurations and dynamics. The possible implications of the mutations have also been discussed, which will provide valuable insights for targeting N protein as an efficient antiviral drug target, vaccine candidate, and development of a more effective diagnostic assays.

## 2. Methodology

### 2.1 Sequence retrieval and processing

To find out the genetic variations of the N protein, we retrieved 67,124 complete (or near-complete) genome sequences of SARS-CoV-2, available at the global initiative on sharing all influenza data (GISAID) (https://www.gisaid.org/) up to July 17, 2020. These sequences belonged to different patients infected with SARS-CoV-2 reported from 145 countries and/or regions of throughout six continents of the world (Supplementary Data 1). Sequences were aligned, and nonsynonymous mutations in the N protein were detected with respect to reference genome from Wuhan (NCBI accession no. NC_045512) using a plug and play pipeline described in previous literature (Rahman et al., 2020). We found 61,485 cleaned sequences of N protein in which 30,221 and 28,327 mutated strains in nucleotide-and amino acid (aa) level, respectively (Supplementary Data 1).

### 2.2 Wu-Kabat protein variability coefficient

To calculate the aa position variability in regards to evolutionary replacements, we employed the Wu-Kabat variability coefficient (Garcia-Boronat et al., 2008; Kabat et al., 1977). The variability coefficient was calculated using the following formula:

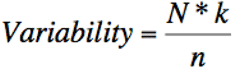

N = total number of sequences in the alignment, k = number of different aa at a given position, n = frequency of the most common aa at that position.

The details calculation of the variability coefficient has been described in Supplementary Data 2.

### 2.3 Tertiary (3D) structure prediction and analyses

(Garcia-Boronat et al., 2008; Kabat et al., 1977)The 3D structure of the N protein of reference genome (NC_045512) was built in Phyre2 (Kelley et al., 2015), followed by energy minimization using the GROMOS96 program (van Gunsteren et al., 1996) implemented within the Swiss-PdbViewer v4.1.0 (Guex and Peitsch, 1997). The 3D structure was validated using the Ramachandran plot (>90 restudies were allowed region) in PROCHECK v3.5 server (https://servicesn.mbi.ucla.edu/PROCHECK/) (Laskowski et al., 1993), and Z-scores (−6.2) from ProSA-web protein structure analysis server (https://prosa.services.came.sbg.ac.at/prosa.php) (Wiederstein and Sippl, 2007). Finally, the energy minimized and validated 3D structure was visualized by PyMol v2.4 (DeLano, 2002). The tertiary structure of N-terminal domain (NTD)/ RNA binding domain (6M3A), and C-terminal domain (CTD)/ dimerization domain (6YUN) available at RCSB Protein Data Bank (Kang et al., 2020; Zinzula et al., 2020) were used for validation of the complete N protein structure built in Phyre2 and observation of the effect of mutations on the domains. The structural NTD and CTD domain positions of newly built N protein were cross checked using the structure of NTD and CTD from protein data bank (Supplementary Fig. 1). The predicted full-length tertiary structure of N protein of the Wuhan strain was used in molecular dynamics study and tracking the deleted aa position of N protein of SARS-CoV-2 in this study.

**Figure 1:**
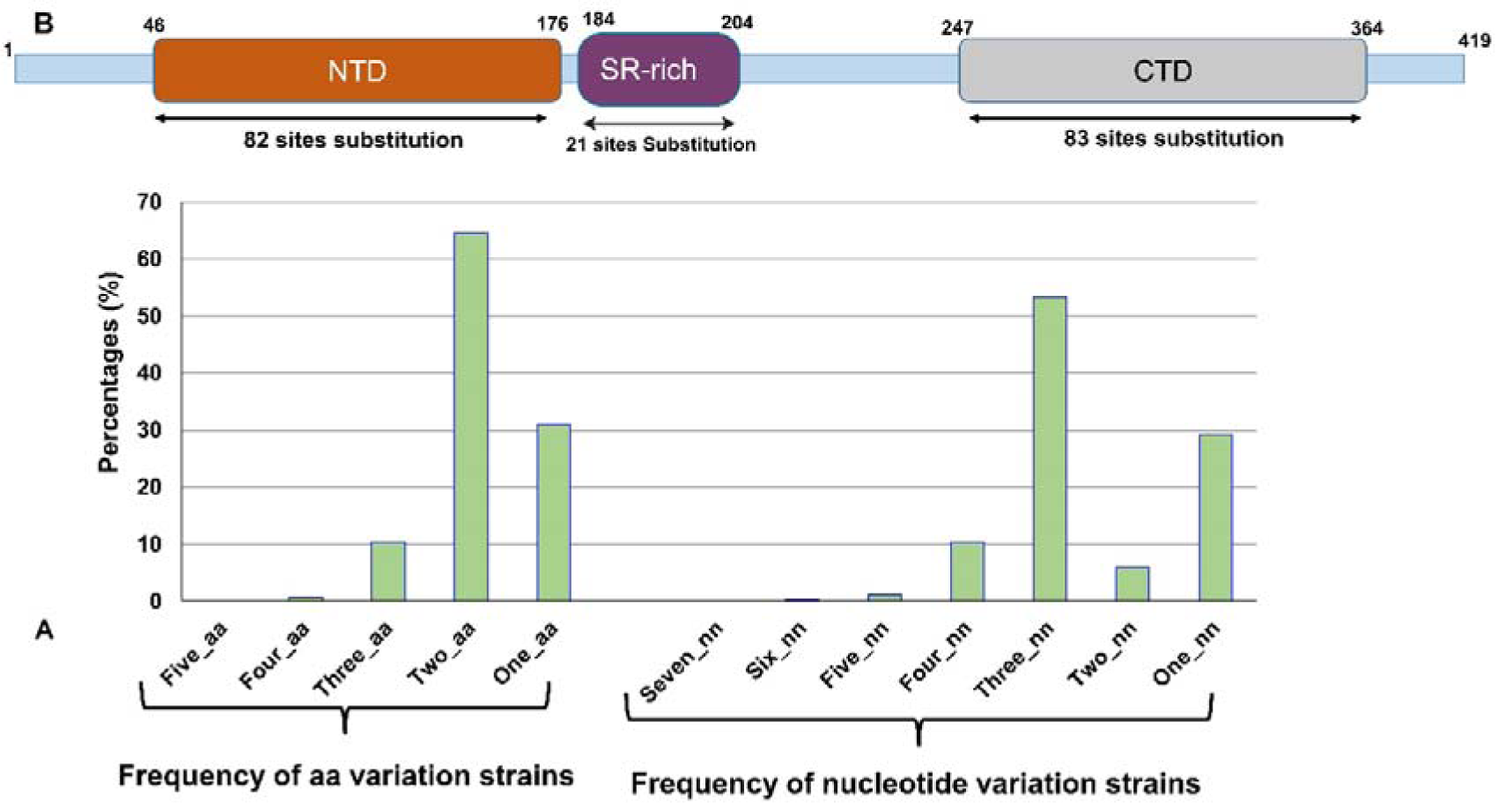
Mutational spectra of SARS-CoV-2 N protein. (A) From aa mutated strains, 64.60 % sequences carried two aa mutations throughout the N protein. The remaining 30.85 %, 10.21 %, 0.67 %, and 0.03 % mutated sequence of N protein had experienced 1, 3, 4, and 5 aa changes, respectively. 53.21 % sequences carried three nucleotide mutation throughout the N and the remaining 29.15 %, 10.30 %, 5.98 %, 1.12 %, 0.20 % and 0.04 % strains showed 1, 4, 2, 5, 6, 7 nucleotide changes, respectively. (B) N-terminal RNA binding domain (NTD) undergoes 82 sites of aa variation, serine/arginine-rich (SR-rich) region 21 aa substitutions, and C-terminal dimerization domain (CTD) undergoes 83 sites of aa mutation.

### 2.4 Consensus building of NTD, CTD domain and their structural analyses

To explore the effect of high frequency mutations in secondary and tertiary structure, we also built consensus for NTD-wild (Wuhan reference) and CTD-wild (Wuhan reference) of the N protein using the top 20 or more mutational frequency. For NTD-cons, we changed 14 aa (A55S, E62V, P67T, D81Y, D103Y, A119S, P122L, D128Y, L139F, D144Y, A152S, A156S, L161F, P168S) while that for CTD-cons, we also changed 13 aa (T247I, A252S, Q289H, I292T, H300Y, T325I, S327L, T334I, D340N, P344S, D348H, T362I, P364L) from the N protein of reference strain. CFSSP (Chou and Fasman secondary structure prediction) was used to predict the secondary structure of NTD-wild, NTD-cons, CTD-wild, and CTD-cons of N protein of the SARS-CoV-2 (Kumar, 2013). Then we built a 3D structure of NTD-cons and CTD-cons through Swiss-Model homology modelling using 6M3A and 6YUN as a template (Waterhouse et al., 2018). Structural differences of NTD-wild and NTD-cons, and CTD-wild and CTD-cons were visualized in PyMol v2.4.

### 2.5 Protein stabilization analysis

To predict the single mutational effect on NTD and CTD protein tertiary structure, molecular stability, and flexibility, we employed DynaMut server (http://biosig.unimelb.edu.au/dynamut/) (Rodrigues et al., 2018). To run this tool, we considered the NTD (6M3A), and CTD (6YUN) structures as a reference, and consequences of the most frequently occurred mutations (NTD-A55S, E62V, P67T, D81Y, D103Y, A119S, A119V, P122L, D128Y, L139F, D144Y, A152S, A156S, L161F, P168S; SR-rich domain-R203K, G204R; CTD-A252S, Q289H, I292T, H300Y, T325I, S327L, T334I, D340N, P344S, D348H, T362I, P364L) were predicted. Protein structure stability parameters such as atomic fluctuations, vibrational entropy, deformation energies, and mutational impact on protein stability were also predicted by the DynaMut server with a well-established predictive normal mode approach. We have calculated free energy difference (ΔΔG) and vibrational entropy energy (ΔΔS_Vib_ ENCoM) by using Dynamut which gives us the report about the impact in protein stability and flexibility in response to mutations in the wild-type structure. Structural changes, such as changes in cavity volume, packing density,, and accessible surface area were correlated with the free energy differences (ΔΔG) and that’s why it is an indicator of the effect of a mutation on protein stability (Eriksson et al., 1992). The ΔΔG value below zero indicates that the mutation causes destabilization and above zero means protein stabilization. For two or more mutational effect prediction, we used FoldX v.5.0 with their default parameters and three times run (Delgado et al., 2019) plugin in YASARA. Measurement of Gibbs free energy (ΔΔG) between mutant and WT (ΔΔG = ΔGmutant – ΔGWT) in unfolding state, FoldX predicts how much protein mutations get into the stability. FoldX first repaired the structure and then predict the mutational effect. The foldX free energy value indicates highly destabilizing (ΔΔG > +1.84 kcal/mol), destabilizing (+0.92 kcal/mol < ΔΔG ≤ +1.84 kcal/mol), slightly destabilizing (+0.46 kcal/mol < ΔΔG ≤ +0.92 kcal/mol), neutral (–0.46 kcal/mol < ΔΔG ≤ +0.46 kcal/mol), slightly stabilizing (–0.92 kcal/mol ≤ ΔΔG < –0.46 kcal/mol), stabilizing (–1.84 kcal/mol ≤ ΔΔG < –0.92 kcal/mol), highly stabilizing (ΔΔG < –1.84 kcal/mol) (Buß et al., 2018; Studer et al., 2014).

## 3. Results and Discussion

### 3.1 Synonymous and nonsynonymous mutations in N protein gene implicates a continuous evolution of SARS-CoV-2

A total of 61,485 cleaned full-length N protein gene sequences were obtained through trimming of the low quality, ambiguous, and non-human host sequences from 67,124 complete (or near-complete) genome sequences of SARS-CoV-2 available in GISAID reported from 145 countries and/or regions of the six continents of the world till 17 July, 2020 (Supplementary Data 1). Compared to the reference genome of the SARS-CoV-2 (Wuhan-Hu-1 strain: Accession NC_045512.2), mutational analyses revealed 49.15 % (n = 30,221) nucleotide-level mutant strains, and 93.73 % (n = 28,327) of the mutant strains carrying at least one aa substitutions in their N protein sequences (Supplementary data 2). Interestingly, 64.60% (n = 17,205) sequences carrying two aa mutations throughout the N protein, whereas a maximum five aa substitutions found in strains of England, USA, and India (Fig. 1A, Supplementary Data<u>2</u>). Moreover, country or region-specific aa mutation patterns revealed the highest percent of N protein mutated SARS-CoV-2 strains in Wales (68.69%) followed by India (68.28 %), England (67.70 %), Scotland (51.29 %), Spain (42.95 %), Australia (37.51 %), and USA (17.87 %) (Supplementary Data 1). However, we did not find any aa substitution in the full-length N protein sequences of 29 countries and/or regions (Supplementary Data 1).

In this study, we found 1034 nucleotide substitutions in 745 unique positions of the N protein gene. Of the nucleotide mutated strains, 53.21% (n = 16,081) sequences carried three nucleotide mutations throughout the N gene while 12 strains from England, India, and USA showed the highest number (n = 7) of nucleotide mutations (Fig. 1, Supplementary Data 2). Overall, these nucleotide changes resulted in 684 nonsynonymous aa substitutions distributed across 317 unique positions of the SARS-CoV-2 N protein (Supplementary Data 2). Therefore, considering the total length of N protein (419 aa), our analysis identified 75.66 % aa positions undergoing evolutionary changes in the SARS-CoV-2 nucleocapsid globally. Simultaneously, 59.27% positions of the nucleotide primary structure (n = 1,257) encoding N-protein showed evolutionary changes (Supplementary Data 2). We also found 391 codon mutation at nucleotide level, of those 81.07% mutations were nonsynonymous. A recent study based on a set of 38,318 genome sequences reported a total of 10,983 instances of aa substitutions and observed change in ∼60% aa positions (250 out of 419 aa residues) in N protein (Ye et al., 2020). These results are implying the continuous evolution of the N protein of SARS-CoV-2.

The two largest domains of N protein, NTD (RNA binding Domain) and CTD (Dimerization domain), had 82 and 83 aa positions substitutions, respectively, on the other hand 21 aa substitutions found within the highly flexible SR-rich domain of the linker region (Fig. 1B). Amino acid group transformation among polar, nonpolar, positively charged, and negatively charged across the NTD and CTD regions may alter the interaction of N protein and viral RNA, and thus affect viral replication, pathogenicity, RNA stability, and disease severity (Surjit and Lal, 2010; Wang et al., 2010). Noteworthy, our previous study found 47.57 % aa positions (606 out of 1274 aa) of the S protein undergoing aa-level evolution worldwide till June 12, 2020 (Rahman et al., 2020). However, some previous reports claimed the N gene more conserved and fewer mutations over time (Dutta et al., 2020; Grifoni et al., 2020). In contrast with these reports, our findings reveal more evolutionary nature with frequent mutations of the N gene compared to the S gene of SARS-CoV-2, which reflects the fragility of the N protein as an alternative vaccine target to the S protein of SARS-CoV-2.

The aa position 180 of the N protein was found to be hypervariable showing 6 aa variations in this particular position. We found 9 sites (aa positions: 13, 80, 185, 191, 195, 203, 216, 378, 385) exhibiting 5 types of aa variations, and 30, 72, 92, and 113 unique sites showed 4, 3, 2, and 1 aa variability, respectively (Supplementary Data 2). One aa position (80) of the NTD showed 5 types aa variations while two aa positions (128, 144) of the NTD showed 4 types of aa variations, followed by 15 and 23 sites showing 3 and 2 types of aa variations, respectively (Table 1). Similarly, the CTD region of the N protein possessed 4, 15 and 27 sites showing 4, 3, and 2 aa variations, respectively (Table 2). The aa variabilities across these domains included the non-polar, polar, positively, or negatively charged groups (Table 1, Table 2). Furthermore, of the five aa variations at position 203 (R203K/M/S/I/G) within the SR-rich domain, the substitution R203K occurred most frequently in 68.09% of the mutated strains globally followed by the substitution G204R found in 67.94% mutated strains of the SARS-CoV-2 (Fig. 2; Supplementary Data 2). Zhou et al. (2020) reported a distinctive polymorphism in SR-rich fragment of SARS-CoV-2 N protein with an extra SRXX repeat and divided this variable segment into 23 different types based on 3233 sequences that alarms of a constant evolution of the SR-rich region (Zhou et al., 2020).

**Table 1:** NTD region aa acid variation and its characteristics of SARS-CoV-2 N protein.

| NTD region aa position | Variations of aa | Name of aa | Reference aa characteristics | aa characteristics |
| --- | --- | --- | --- | --- |
| 80 | 5 | P80L,P80R,P80S,P80Q,P80T | NP | NP,PC,P,P,P |
| 128 | 4 | D128Y,D128H,D128E,D128N | NC | P,PC,NC,P |
| 144 | 4 | D144Y,D144N,D144H,D144E | NC | P,P,PC,NC |
| 62 | 3 | E62V,E62K,E62G | NC | NP,PC,NP |
| 63 | 3 | D63Y,D63N,D63H | NC | P,P,PC |
| 67 | 3 | P67T,P67S,P67L | NP | P,P,NP |
| 68 | 3 | R68Q,R68L,R68G | PC | P,NP,NP |
| 70 | 3 | Q70R,Q70L,Q70H | P | PC,NP,PC |
| 72 | 3 | V72A,V72I,V72L | NP | NP,NP,NP |
| 103 | 3 | D103F,D103Y,D103N | NC | NP,P,P |
| 119 | 3 | A119V,A119S,A119P | NP | NP,P,NP |
| 120 | 3 | G120V,G120R,G120A | NP | NP,PC,NP |
| 135 | 3 | T135I,T135N,T135P | P | NP,P,NP |
| 141 | 3 | T141I,T141P,T141A | P | NP,NP,NP |
| 145 | 3 | H145Q,H145Y,H145N | PC | P,P,P |
| 151 | 3 | P151L,P151H,P151S | NP | NP,PC,P |
| 161 | 3 | L161H,L161F,L161P | NP | PC,NP,NP |
| 168 | 3 | P168S,P168Q,P168L | NP | P,P,NP |
| 50 | 2 | A50E,A50V | NP | NC,NP |
| 61 | 2 | K61R,K61N | PC | PC,P |
| 65 | 2 | K65E,K65I | PC | NC,NP |
| 81 | 2 | D81Y,D81H | NC | P,PC |
| 90 | 2 | A90S,A90T | NP | P,P |
| 95 | 2 | R95C,R95L | PC | NP,NP |
| 101 | 2 | M101I,M101L | NP | NP,NP |
| 122 | 2 | P122L,P122S | NP | NP,P |
| 125 | 2 | A125S,A125T | NP | P,P |
| 129 | 2 | G129S,G129D | NP | P,NC |
| 134 | 2 | A134V,A134S | NP | NP,P |
| 139 | 2 | L139F,L139V | NP | NP,NP |
| 140 | 2 | N140T,N140K | P | P,PC |
| 142 | 2 | P142S,P142L | NP | P,NP |
| 146 | 2 | I146T,I146F | NP | P,NP |
| 152 | 2 | A152S,A152V | NP | P,NP |
| 157 | 2 | I157T,I157V | NP | P,NP |
| 162 | 2 | P162L,P162S | NP | NP,P |
| 163 | 2 | Q163R,Q163K | P | PC,PC |
| 164 | 2 | G164R,G164V | NP | PC,NP |
| 165 | 2 | T165I,T165P | P | NP,NP |
| 170 | 2 | G170S,G170C | NP | P,NP |
| 173 | 2 | A173V,A173T | NP | NP,P |

\*NP=Non-Polar, Charge, P=Polar, PC=Positive, NC=Negative Charge

**Table 2:** CTD region aa acid variation and its characteristics of SARS-CoV-2 N protein.

| CTD region aa position | Variations of aa | Name of aa | Reference aa characteristics | aa characteristics |
| --- | --- | --- | --- | --- |
| 252 | 4 | A252S,A252D,A252V,A252T | NP | P,NC,NP,P |
| 253 | 4 | E253A,E253G,E253V,E253D | NC | NP,NP,NP,NC |
| 289 | 4 | Q289H,Q289L,Q289R,Q289K | P | PC,NP,PC,PC |
| 322 | 4 | M322I,M322T,M322V,M322L | NP | NP,P,NP,NP |
| 254 | 3 | A254T,A254S,A254V | NP | P,P,NP |
| 255 | 3 | S255F,S255A,S255P | P | NP,NP,NP |
| 272 | 3 | Q272L,Q272R,Q272H | P | NP,PC,PC |
| 281 | 3 | Q281L,Q281R,Q281K | P | NP,PC,PC |
| 292 | 3 | I292T,I292V,I292L | NP | P,NP,NP |
| 310 | 3 | S310I,S310C,S310N | P | NP,NP,P |
| 321 | 3 | G321D,G321R,G321C | NP | NC,PC,NP |
| 325 | 3 | T325I,T325R,T325A | P | NP,PC,NP |
| 326 | 3 | P326S,P326L,P326H | NP | P,NP,PC |
| 340 | 3 | D340N,D340G,D340Y | NC | P,NP,P |
| 343 | 3 | D343Y,D343V,D343G | NC | P,NP,NP |
| 348 | 3 | D348H,D348Y,D348N | NC | PC,P,P |
| 351 | 3 | I351V,I351T,I351M | NP | NP,P,NP |
| 363 | 3 | F363L,F363S,F363V | NP | NP,P,NP |
| 364 | 3 | P364S,P364L,P364R | NP | P,NP,PC |
| 247 | 2 | T247I,T247A | P | NP,NP |
| 248 | 2 | K248N,K248R | PC | P,PC |
| 250 | 2 | S250F,S250P | P | NP,NP |
| 251 | 2 | A251V,A251S | NP | NP,P |
| 263 | 2 | T263I,T263A | P | NP,NP |
| 265 | 2 | T265I,T265A | P | NP,NP |
| 270 | 2 | V270L,V270I | NP | NP,NP |
| 271 | 2 | T271I,T271A | P | NP,NP |
| 279 | 2 | P279L,P279S | NP | NP,P |
| 283 | 2 | Q283H,Q283K | P | PC,PC |
| 290 | 2 | E290D,E290Q | NC | NC,P |
| 294 | 2 | Q294L,Q294K | P | NP,PC |
| 300 | 2 | H300Y,H300Q | PC | P,P |
| 306 | 2 | Q306L,Q306H | P | NP,PC |
| 311 | 2 | A311V,A311S | NP | NP,P |
| 319 | 2 | R319H,R319S | PC | PC,P |
| 324 | 2 | V324I,V324F | NP | NP,NP |
| 327 | 2 | S327L,S327P | P | NP,NP |
| 329 | 2 | T329M,T329A | P | NP,NP |
| 334 | 2 | T334S,T334I | P | P,NP |
| 336 | 2 | A336S,A336V | NP | P,NP |
| 337 | 2 | I337V,I337F | NP | NP,NP |
| 344 | 2 | P344L,P344S | NP | NP,P |
| 347 | 2 | K347R,K347N | PC | PC,P |
| 350 | 2 | V350F,V350A | NP | NP,NP |
| 359 | 2 | A359T,A359S | NP | P,P |
| 362 | 2 | T362I,T362K | P | NP,PC |

\*NP=Non-Polar, Charge, P=Polar, PC=Positive, NC=Negative Charge

**Figure 2:**
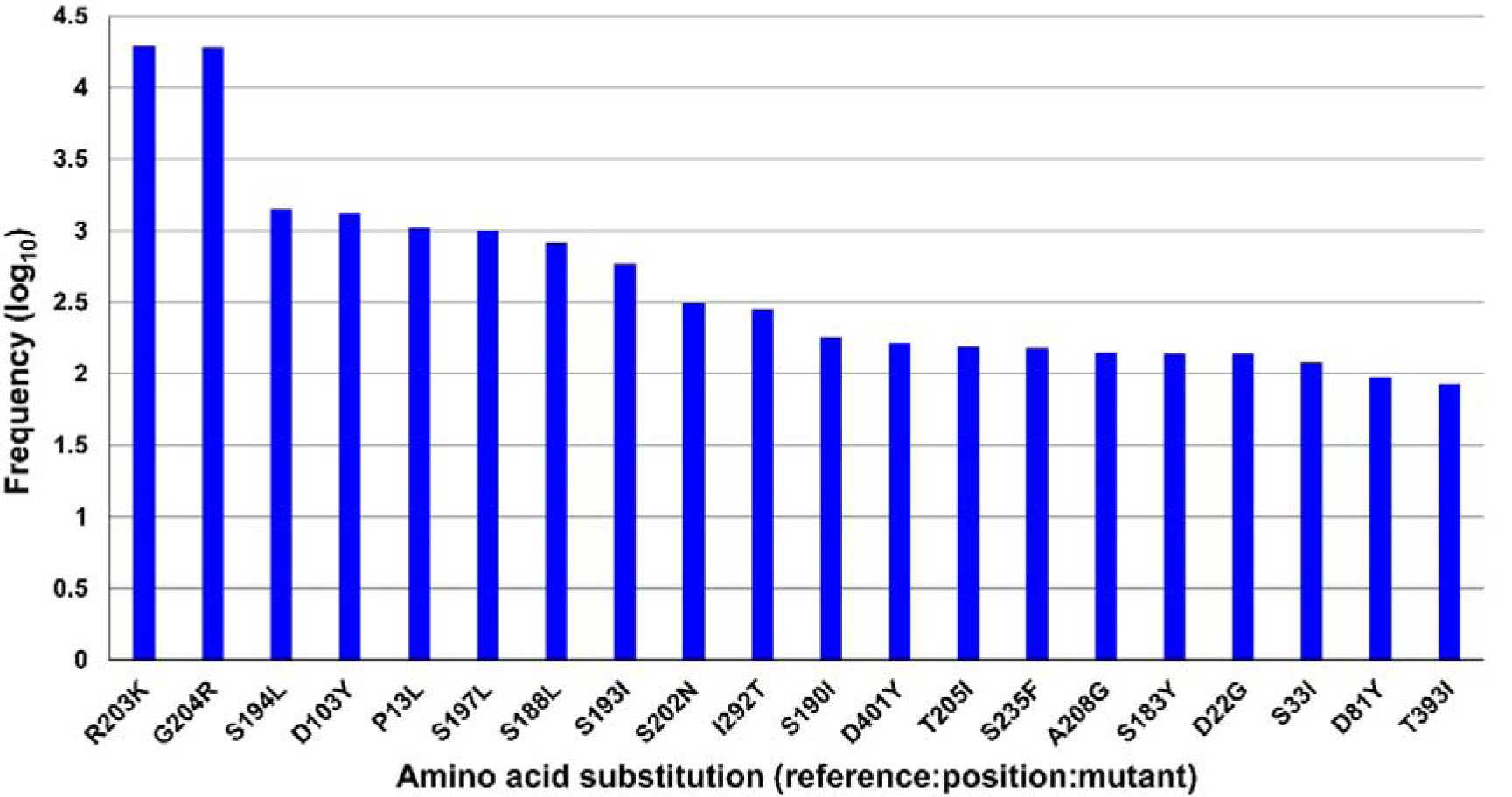
Top 20 high-frequency mutations in SARS-CoV-2 N protein.

It is noteworthy that the NTD and CTD regions are predicted to encompass major antigenic sites of the N protein of the SARS-CoV (Surjit and Lal, 2009). Antibodies against the N protein could persist for a longer period, and may occur in greater abundance in SARS-CoV patients than antibodies against other structural proteins such as the membrane, spike,, and envelope proteins (Wang et al., 2010). This could be due to the expression of higher levels of N protein, compared with other viral proteins, after SARS-CoV-2 infection. Similarly, SARS-CoV-2 N protein is strongly antigenic, and thus could play a significant role in the of the host immune response generation, and preventing the immunopathological damage (Dutta et al., 2020). Thus, aa variations in the antigenically critical domains of N protein may alter immunopathogenicity of the virus, which needs to be considered before the therapeutic interventions of the N protein.

The Wu-Kabat protein variability plot showed the position-specific aa variations with respect to their frequency (Fig. 3). The current variability analysis identified 41 positions showing Wu-Kabat variability coefficient [5, where coefficient 1 indicates no variability compared to the reference strain. The maximum coefficient found for position 203 (coefficient, 8.75) followed by positions 180 (7.01), 13 (6.11), 195 (6.01), 185 (6.01), 191 (6.01), 378 (6.01), 80 (6.00), 216 (6.00), and 385 (6.00) (Fig. 3, Supplementary Data 2). Of the structural aa (n = 419) of the N protein, 24.10 % of positions had protein variability coefficient 1 indicating the conservancy of these positions.

**Figure 3:**
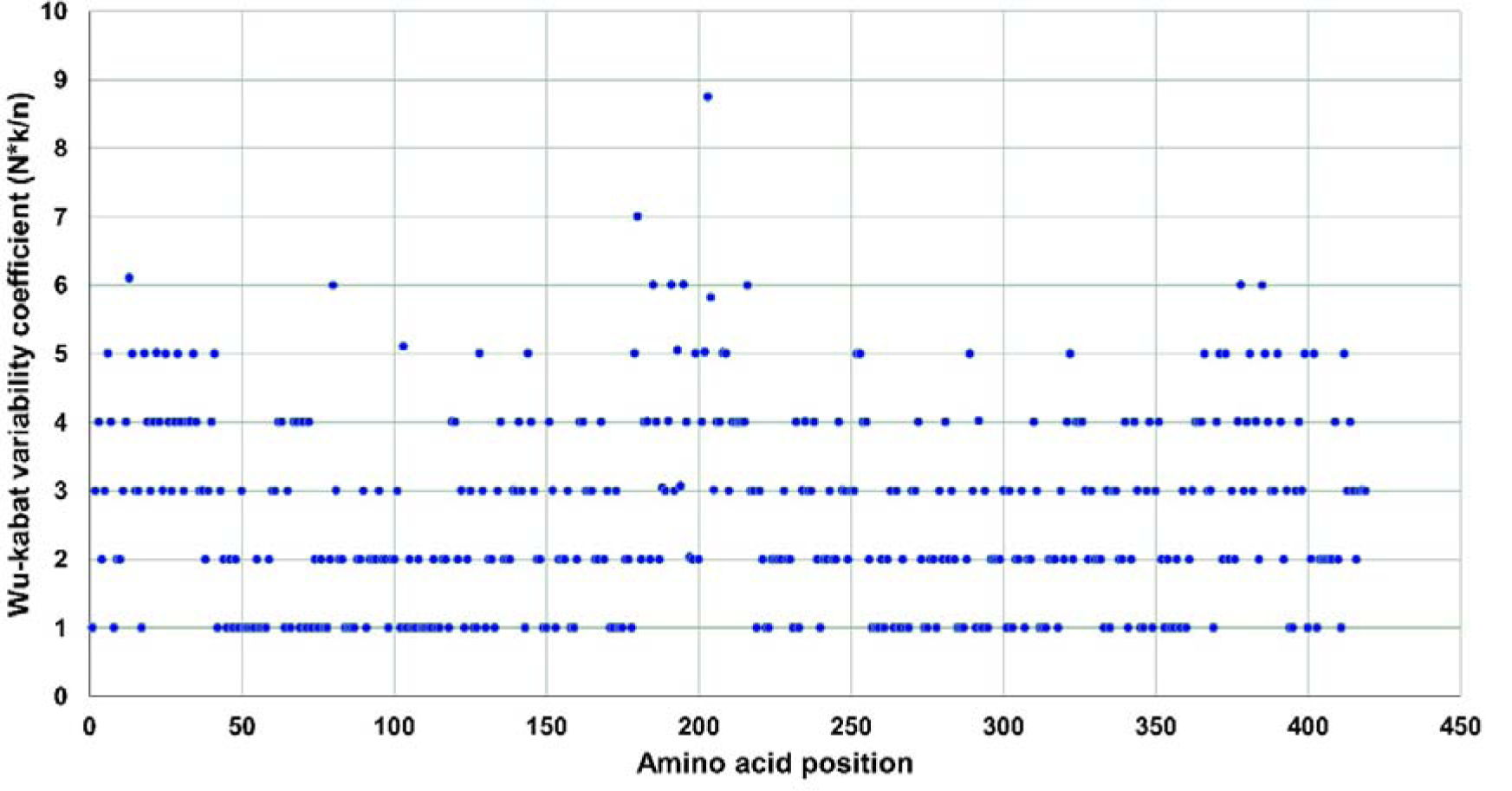
Wu-kabat protein variability coefficient plot of SARS-CoV-2 N protein. Wu-kabat variability coefficient 1 indicates no variability in that position. Highest aa variability found in position 203.

### 3.2 Frequent nonsynonymous mutations impact upon N protein structure and stability

To observe the possible impact on secondary and tertiary structure of the NTD and CTD domain of N protein of these frequent nonsynonymous mutations, secondary and tertiary structures were analyzed. It revealed changes in helix, sheet, and turn composition in the secondary structure of the consensus (Fig. 4). These findings indicated the significant impact of the high –frequency nonsynonymous substitutions on the secondary structure of NTD and CTD domains of the SARS-CoV-2 nucleocapsid, and changing the secondary structure may alter the function of these domains. Furthermore, the homology consensus modeling of this study revealed that the substitutions causing aa group transversions affected the overall tertiary structure of the domains (Fig. 5, Table 1, Table 2). Group changing aa may significantly affect the ultimate functions of the N protein including virus replication, pathogenesis, and disease severity (Islam et al., 2020b). Furthermore, the higher mutation rates of this protein in India, Wales, and European countries warrants the effect of the geo-climatic conditions, and ethnic groups. The current evolutionary findings offer important and timely insight relevant to the SARS–CoV-2 N protein heterogeneity throughout the world which could be associated with virus pathogenesis and disease severity.

**Figure 4:**
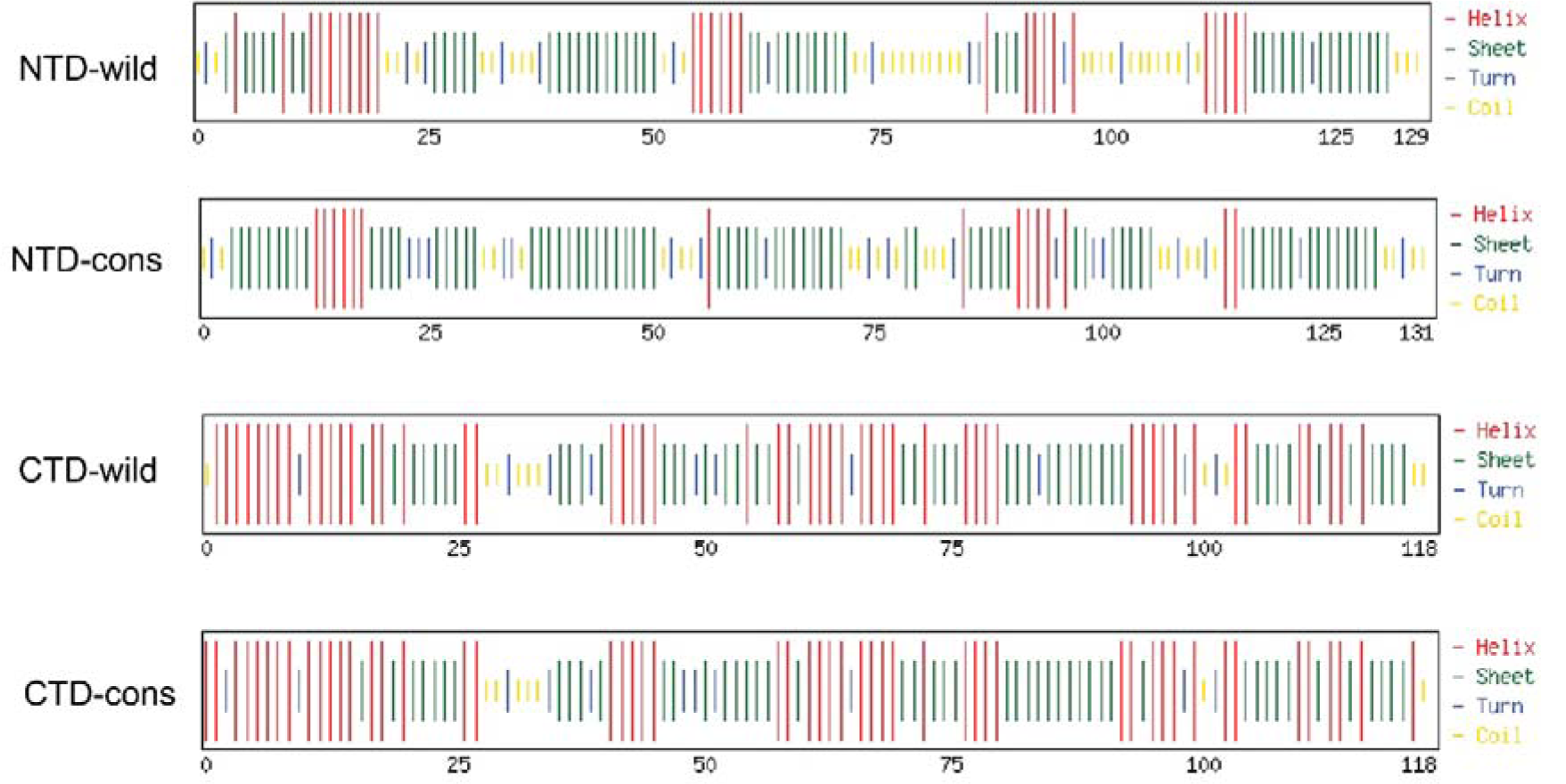
Chou and Fasman secondary structure prediction of NTD and CTD domain of SARS-CoV-2 N protein. For NTD-wild type, there are Helix 36.4 %, Sheet 45.0 %, Turn 15 %; on the other hand, NTD-consensus type, there are Helix 28.2 %, Sheet 64.1 %, Turn 16 %. For CTD-wild type, there are Helix 71.2 %, Sheet 68.6 %, Turn 16.1 %; whereas in CTD-consensus we found Helix 80.5 %, Sheet 64.4 %, Turn 13.6 %.

**Figure 5:**
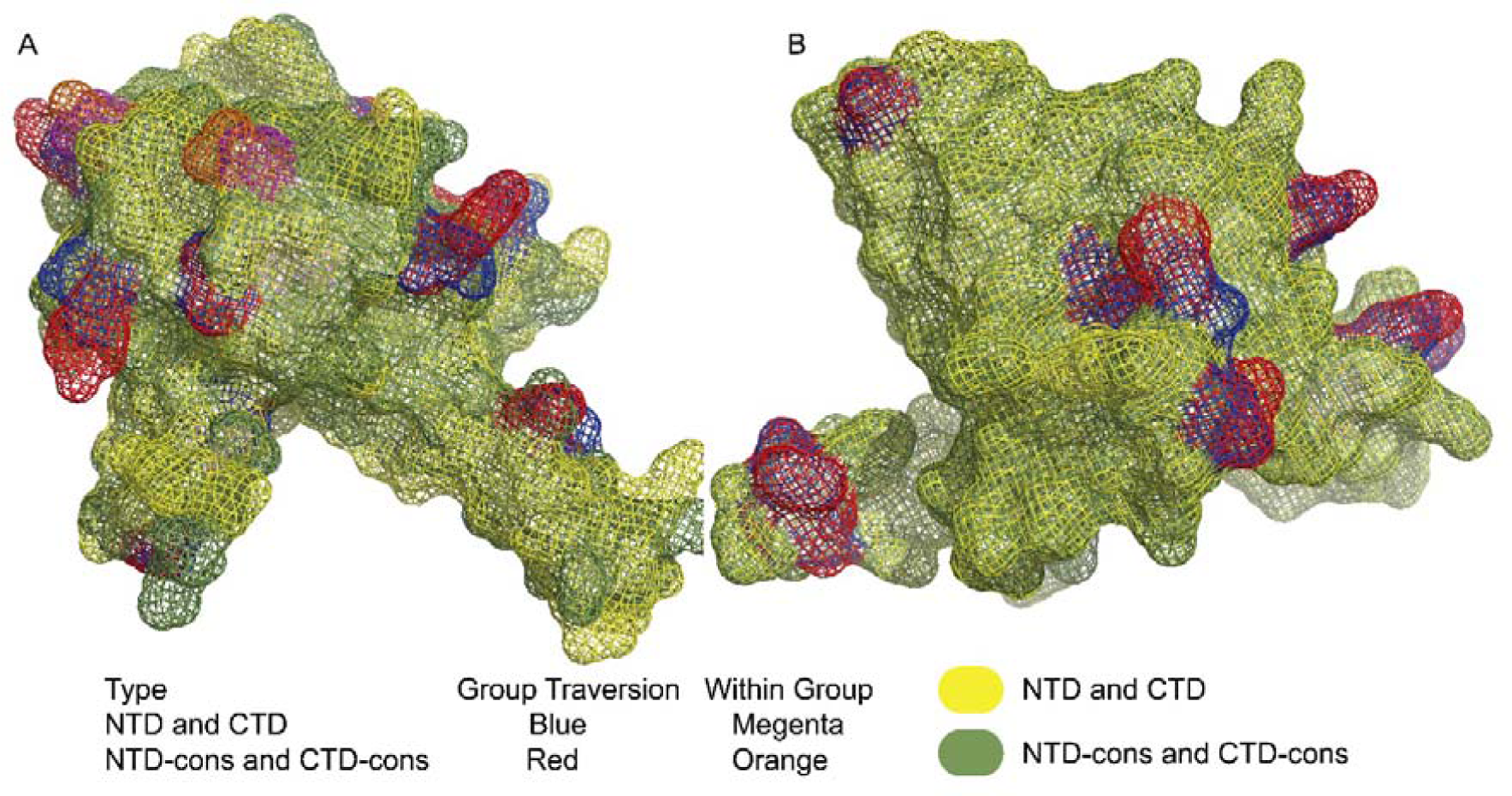
aa transverse effect of NTD and CTD 3D structure. (A) Structural alignment of NTD and NTD-cons. Here from 14 aa, eleven aa (A55S, E62V, P67T, D81Y, D103Y, A119S, D128Y, D144Y, A152S, A156S, P168S) substitutions was in group transversion and rest of the three aa (P122L L139F, L161F) substitutions was within group. (B) Structural alignment of CTD and CTD-cons. Here, from 13 aa, eleven aa (A252S, Q289H, I292T, H300Y, T325I, S327L, T334I, D340N, P344S, D348H, T362I) substations was in group transversion and the other two aa (T247I, P364L) transverse within group. Structures were visualized in PyMol v2.4.

The ΔΔG values for the two high frequency aa substitutions R203K and G204R in the SR-rich region, were 0.460 kcal/mol and 1.361 kcal/mol, respectively implicating these changes stabilize the N structure (modelled in Phyre2). We also closely inspected the changes in the intramolecular interactions due to these two mutations (R203K and G204R) in N protein and found significant variation ((Fig. 6A-D). An average of the configurational entropies of the protein is revealed by vibrational entropy within single minima of the energy landscape (Goethe et al., 2015). For both mutations (R203K, and G204R), the vibrational entropy energy (ΔΔS_Vib_ ENCoM) between the wild-type, and the mutant, ΔΔS_Vib_ ENCoM values were -0.137 kcal.mol^-1^.K^-1^, and -1.497 kcal.mol^-1^.K^-1^ respectively, which indicate the decrease of molecular flexibility in the N protein.

**Figure 6:**
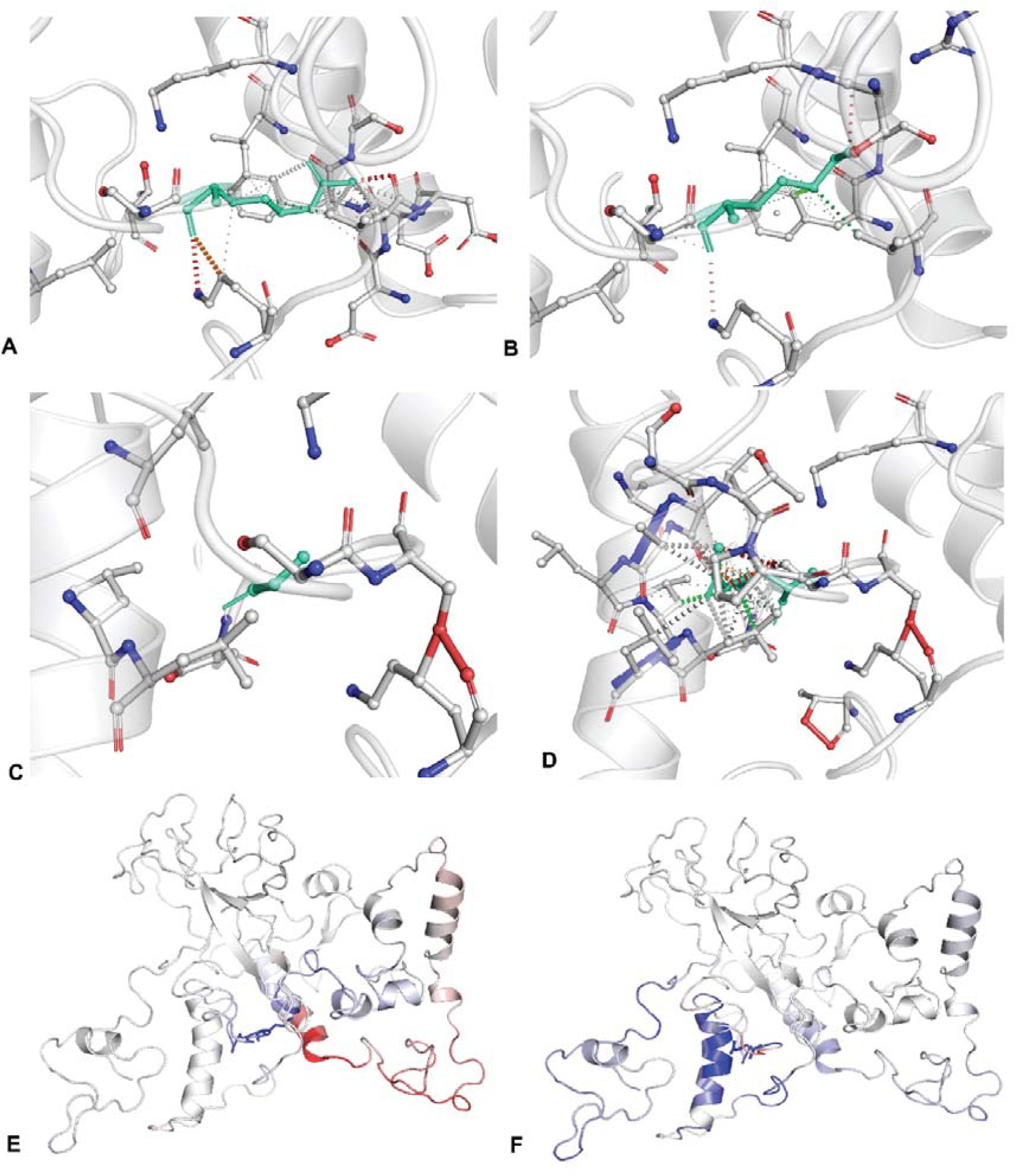
Prediction of Interactomic Interactions and flexibility in response to substitutions. (R203K, G204R). Wild-type and mutant residues are colored in light-green and are also represented as sticks alongside with the surrounding residues which are involved on any type of interactions. Amino acids which are colored according to the vibrational entropy change upon mutation. BLUE represents a rigidification of the structure and RED a gain in flexibility. (A, C) represents the wild type interactomic interaction at position of 203, and 204 respectively and (B, D) represents the mutant interactomic interaction at the same positions, respectively. (E) Overall decrease the flexibility due to R203K mutation and aa positions (201-209, 254-261) gain rigidification whereas aa positions (362-404) gain flexibility. (F) G204R mutation also causes the overall decrease of flexibility and aa positions (21-46, 202-206,215-226, 240-248) gain rigidification but 200-202 aa positions gain flexibility. Structures were visualized in PyMol v2.4.

The overall decrease in the flexibility due to substitution R203K conferred the rigidification of the protein structure at aa positions (201-209, 254-261) whereas that mutation moved to flexibility at aa positions (362-404) (Fig. 6E). The G204R mutation also influenced the overall decrease of flexibility, and thus aa positions (21-46, 202-206,215-226, 240-248) gained rigidification. However, the 200-202 aa positions of the N protein showed optimum flexibility due to G204R mutation (Fig. 6F). Therefore, only the R203K mutation affected the CTD (dimerization domain) at 254-261 aa positions which decreased the flexibility of the domain (Fig. 6E). In our study, we found total strains 19,246 where both R203K and G204R mutation observed throughout the 89 countries or region (Supplementary Data 2).

We have also observed the effect of the double mutations in FoldX (Schymkowitz et al., 2005). For the paired mutations, we found ΔΔG = 3.42262 kcal/mol which indicates highly destabilizing the structure. We also found deletion at the same position in the N protein of SARS-Cov-2 (Table 5). For NTD and CTD structural effect analyses, we considered original structures from RCSB Protein Data Bank 6M3A and 6YUN, respectively. The effects of different high frequency mutation on NTD and CTD are described in Table 3. A55S, P67T, D81Y, A119V, P122L, D128Y, L139F, D144Y substitutions increase the stability of NTD structure whereas E62V, D103Y, A119S, A152S, A156S, L161F, P168S substitutions decrease the stability of that same structure. Again, E62V, P67T, D144Y, P168S substitutions increase the molecular flexibility but A55S, D81Y, D103Y, A119S, A119V, P122L, D128Y, L139F, A152S, A156S, L161F decrease the molecular flexibility of RNA binding domain (Table 3).

**Table 3:** NTD and CTD mutations and their effects on three dimensional structure in Dynamut. (ΔΔG, free energy difference and ΔΔS_Vib_ ENCoM, Vibrational Entropy Energy Between Wild-Type and Mutant).

| aa substitutions | aa characteristics | Frequency of strains | $\Delta\Delta G$ DynaMut (kcal/mol) | Outcome | $\Delta\Delta S_{\text{vib}}$ ENCoM (kcal.mol<sup>-1</sup>.K<sup>-1</sup>) | Molecule flexibility |
| --- | --- | --- | --- | --- | --- | --- |
| **NTD Mutation** |  |  |  |  |  |  |
| A55S | NP to P | 44 | 1.117 | Stabilizing | -0.337 | Decrease |
| E62V | NC to NP | 40 | -0.006 | Destabilizing | 0.392 | Increase |
| P67T | NP to P | 21 | 0.289 | Stabilizing | 0.011 | Increase |
| D81Y | NC to P | 94 | 0.580 | Stabilizing | -0.400 | Decrease |
| D103Y | NC to P | 1324 | -0.126 | Destabilizing | -0.083 | Decrease |
| A119S | NP to P | 78 | -0.114 | Destabilizing | -0.153 | Decrease |
| A119V | NP to NP | 31 | 0.120 | Stabilizing | -0.063 | Decrease |
| P122L | NP to NP | 22 | 0.006 | Stabilizing | -0.126 | Decrease |
| D128Y | NC to P | 60 | 0.654 | Stabilizing | -0.294 | Decrease |
| L139F | NP to NP | 32 | 0.248 | Stabilizing | -0.205 | Decrease |
| D144Y | NC to P | 36 | 0.268 | Stabilizing | 0.027 | Increase |
| A152S | NP to P | 23 | -0.220 | Destabilizing | -0.043 | Decrease |
| A156S | NP to P | 76 | 0.252 | Destabilizing | -0.402 | Decrease |
| L161F | NP to NP | 22 | 0.502 | Destabilizing | -0.255 | Decrease |
| P168S | NP to P | 40 | -0.438 | Destabilizing | 0.150 | Increase |
| **CTD Mutation** |  |  |  |  |  |  |
| A252S | NP to P | 26 | -0.054 | Destabilizing | -0.078 | Decrease |
| Q289H | P to PC | 24 | -0.411 | Destabilizing | 0.095 | Increase |
| I292T | NP to P | 284 | -1.736 | Destabilizing | 0.183 | Increase |
| H300Y | PC to P | 29 | 0.534 | Stabilizing | -0.184 | Decrease |
| T325I | P to NP | 24 | 1.017 | Stabilizing | -0.170 | Decrease |
| S327L | P to NP | 37 | 0.506 | Stabilizing | -0.175 | Decrease |
| T334I | P to NP | 26 | 0.216 | Stabilizing | -0.225 | Decrease |
| D340N | NC to P | 29 | 0.369 | Stabilizing | -0.080 | Decrease |
| P344S | NP to P | 23 | -0.051 | Destabilizing | -0.079 | Decrease |
| D348H | NC to PC | 30 | -0.128 | Destabilizing | -0.040 | Decrease |
| T362I | P to NP | 44 | 0.225 | Stabilizing | 0.054 | Increase |
| P364L | NP to NP | 35 | 0.172 | Stabilizing | 0.108 | Increase |

\*NP=Non-Polar, Charge, P=Polar, PC=Positive, NC=Negative Charge

For CTD domain, H300Y, T325I, S327L, T334I, D340N, T362I, P364L changes increased stability, on the other hand, A252S, Q289H, I292T, P344S, D348H changes decrease molecular stability. T362I, P364L, Q289H, I292T substitutions increase flexibility but A252S, H300Y, T325I, S327L, T334I, D340N, P344S, D348H decrease molecular flexibility (Table 3). We also found that coevolving nonsynonymous mutations (A55S, D103Y) at England (35 strains) and Wales (1 strain); (R93L, D103Y) at England (8 strains); (D103Y, A119P) at England (4 strains); (D103Y, T165I) at England (2 strains); (G97S, N140T) at Russia (4 strains) in NTD region and (271I, T325I) at England (5 strains); (A252S, S310C) at wales (3 strains) in CTD domain of SARS-CoV-2 N protein (Supplementary Data 2). We analyzed the impact of these coevolving nonsynonymous mutations on NTD structure (6M3A) and CTD structure (6YUN) by using FoldX. Coevolving mutations (D103Y, T165I) have stabilizing (ΔΔG = -1.083 kcal/mol) effect, whereas (G97S, N140T) and (R93L, D103Y) substitutions (ΔΔG = -0.739 kcal/mol; ΔΔG = -0.706 kcal/mol) showed slightly stabilizing impact. On the other hand, (D103Y, A119P) mutation have neutral effect (ΔΔG = -0.046 kcal/mol), whereas (A55S, D103Y) substitutions destabilize (ΔΔG = 1.171 kcal/mol) the RNA binding domain.

In dimerization domain (CTD), (T271I, T325I) change have neutral effect (ΔΔG = -0.268 kcal/mol) but A252S and S310C substitutions highly destabilized (ΔΔG = -2.149 kcal/mol) the original CTD structure. Ye at al. 2020 identified RNA-binding domain residues by NMR peak shifts (A50, T57, H59, R92, I94, S105, R107, R149, and Y172). Our analyses observed aa substitutions in RNA-binding domain (6 residues A50E (Wuhan), A50V (England and India), H59P (Morocco), R92S (India), I94V (England), S105N (USA). Noteworthy, a previous study reported only one mutation (A50E) in the putative RNA binding surface (Ye et al., 2020). From these findings, it can be conclusively stated that the high-frequency aa substitutions are changing the stability, and intramolecular interactions of N protein with in the SARS-CoV-2 RNA implicating a critical functional consequence.

### 3.3 Mutations in the primer binding sites might hamper the efficiency of RT-PCR

In our present study, we observed mutational changes in the regions of 11 primer-probe sets recommended for nucleic acid-based detection of SARS-CoV-2 (Table 4). We found a total of 367 mutations within the primer binding regions for all the well characterized primer sets of the studied here (Table 4). The dynamics of mutations in the primer binding region revealed a total of 367 mutations within the primer binding sites of the primer sets studied (Table 4). The primers sets carried 11-31 mutations within forward/reverse primer binding sites. Minimum mismatch (n=11) in forward and reverse primers of CDC (USA) recommended set N2 and forward primers of Hong Kong University and Park et al. (2020), whereas maximum (n=31) mismatches were found in the forward primer binding sites of China, CDC primer set (Table 4). The CDC, USA recommended 3 primer sets showing comparatively fewer mutations than many of the other primer sets (Table 4). Remarkably, no strains found to have 3’ end mismatch with the N1 forward primer (CDC, US), whereas reverse primer of this set showed 27 mutations worldwide including 3’-end mismatch with some strains from Netherlands. SARS-CoV-2 strains from Finland contained 3’-end mismatch with the N2 forward primer (USA, CDC) and the 3’-end of this reverse primer mismatched with some viruses from USA and England. Strains of Senegal and India were found to have 3’-end mismatches with the forward and reverse primers, respectively, in reference to the CDC (USA) recommended N3 primer set. However, taking only 180 strains into consideration, Nalla et al. (2020) found no mismatch within CDC (USA) recommended N2 primer set showing more sensitivity than N1 and N3. In our study, we found comparatively lower mutations (11 mutations individually in forward and reverse primer) in the N2 set, supporting its superiority for COVID-19 detection (Table 4). The Chinese CDC recommended primer set had 3’-end mismatch with the SARS-Cov-2 strains of Luxembourg (forward primer), and India, Canada, Wales, England (reverse primer). Again, forward primers of Hong Kong University and P1 set of Park et al. (2020) had no 3’-end mismatch with the global sequence dataset used in this study (Table 4). However, mutations in the primer binding sites, importantly at 3’ end, may affect the RT-PCR-based COVID-19 detection resulting in false negative results (Nalla et al., 2020; Rana and Pokhrel, 2020).

**Table 4:**
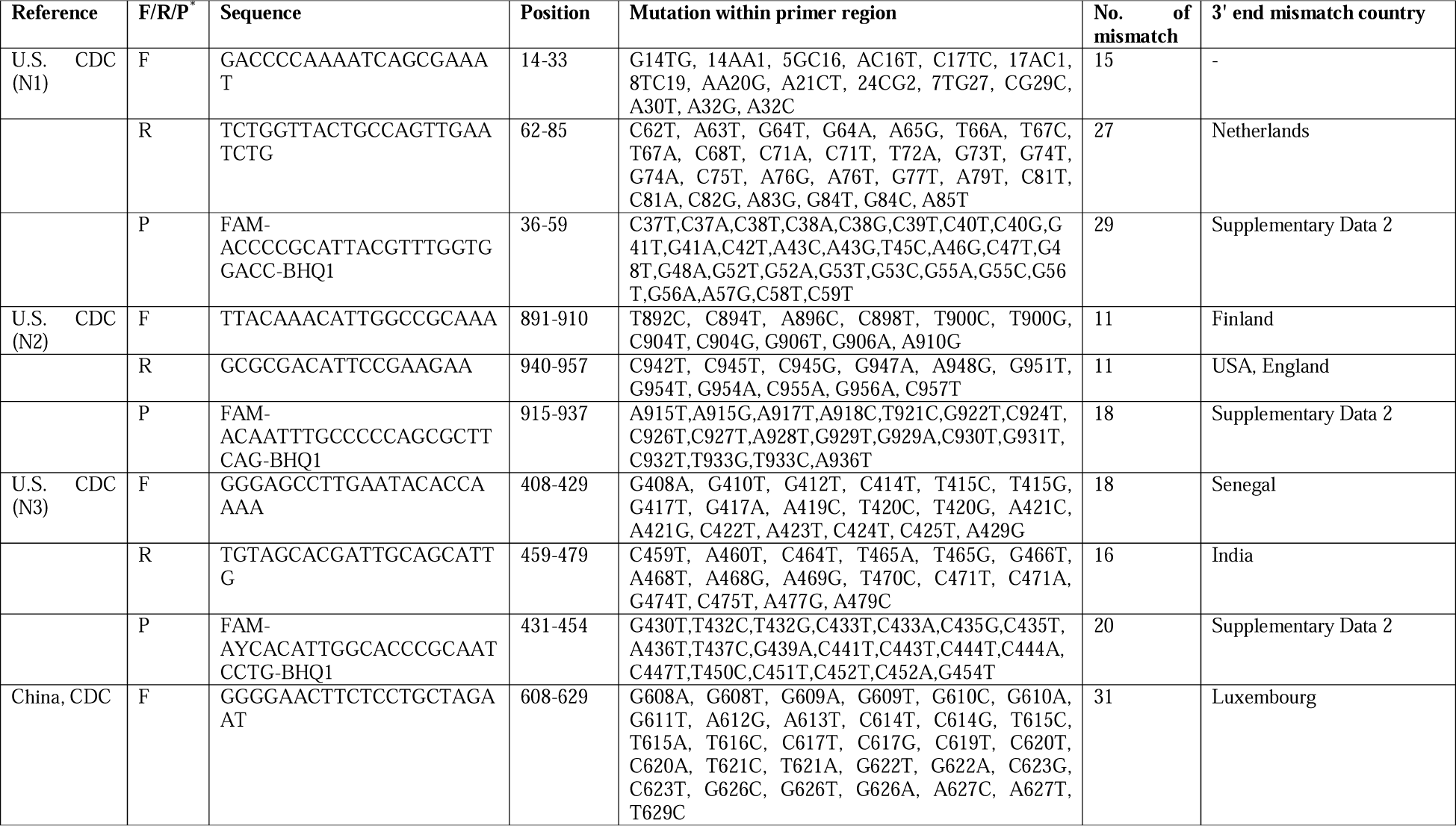
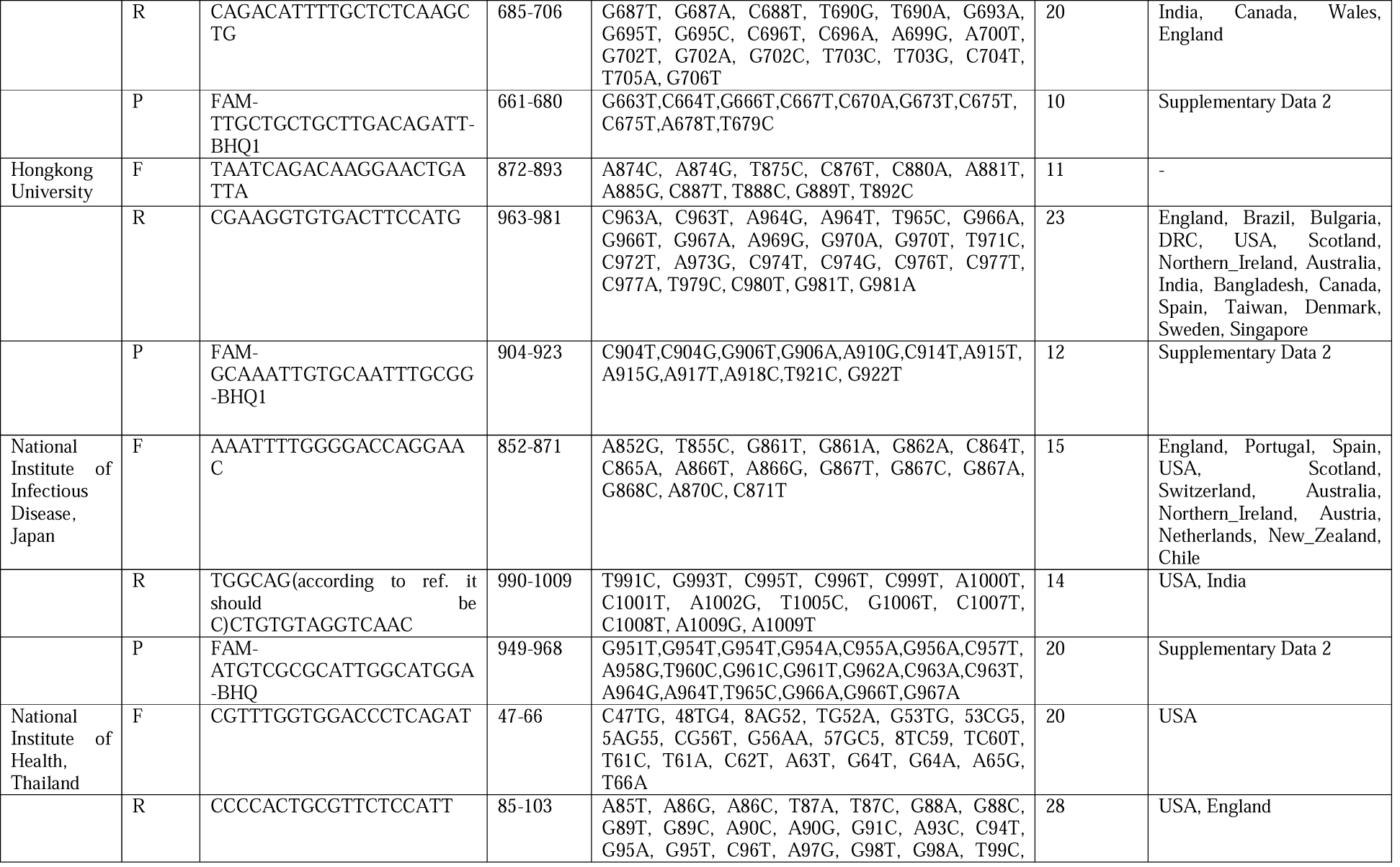
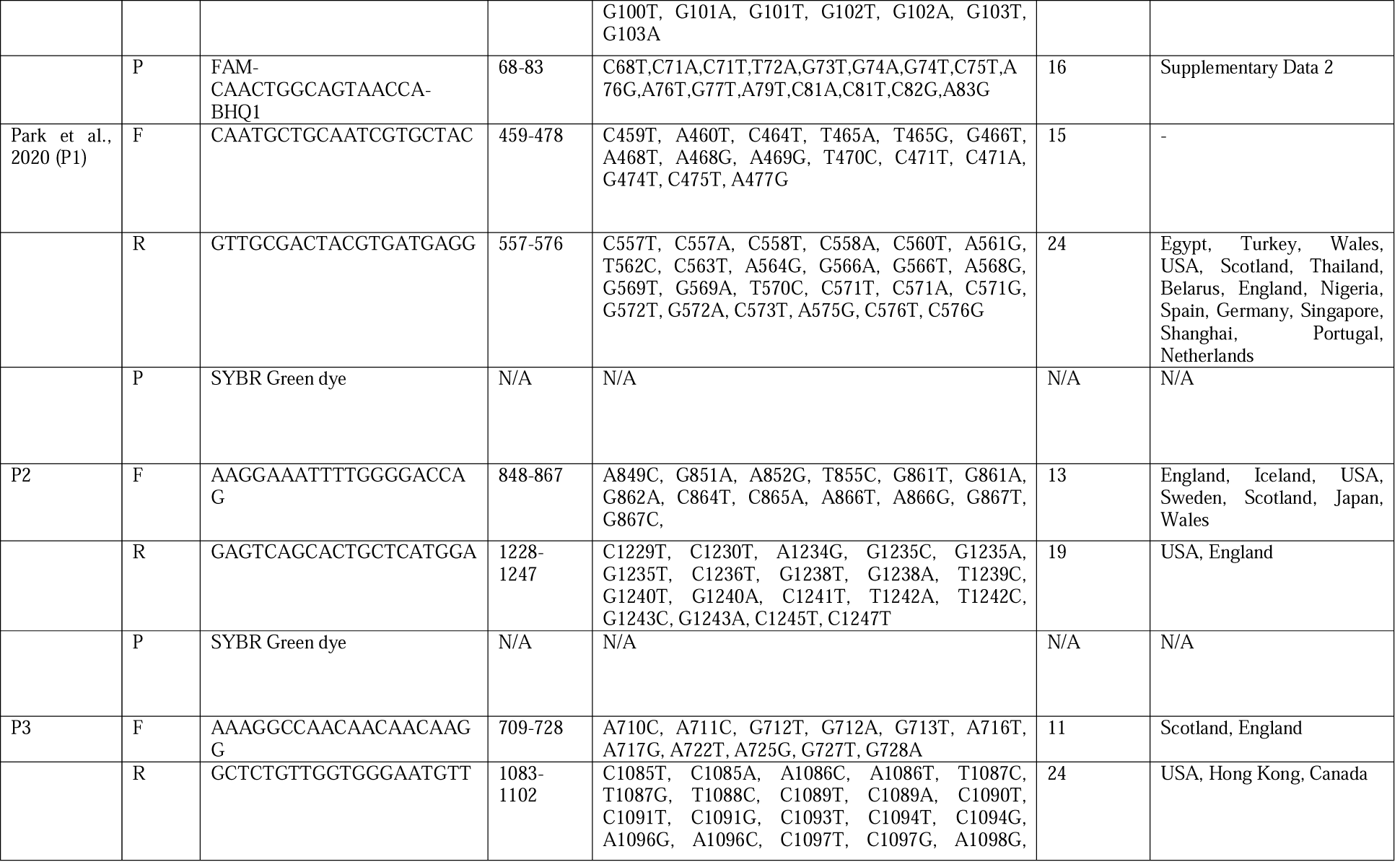
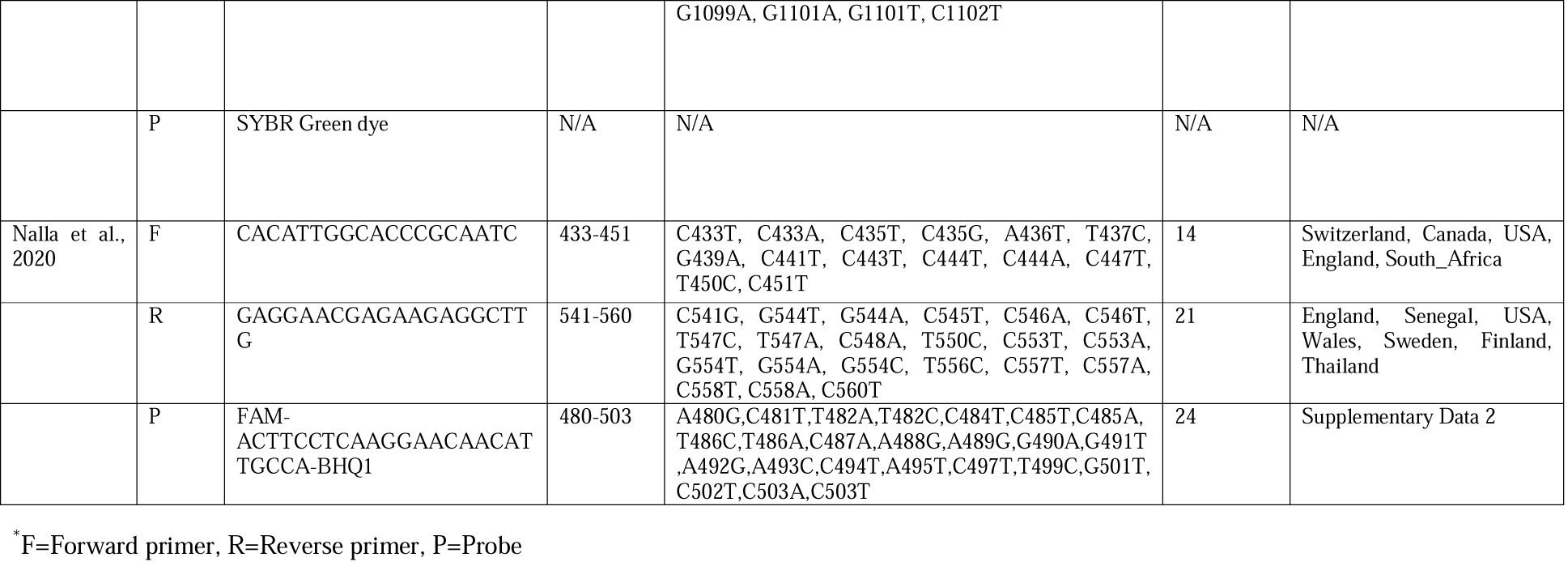
Mutation in primer sequences targeting N gene of SARS-CoV-2 for PCR-based diagnosis.

Moreover, we found notable sequence variations (n = 140) at probe recognition sites of the primers studied here in the virus strains from different countries and/or regions (Table 4, Supplementary data 2). Remarkably, CDC (China) recommended probe showed lowest mismatch (n = 10), whereas probe of CDC (USA) designed N1 primer exhibited maximum variations (n = 29). Other primer-probes carried 12-24 mismatches along the probe recognition site in the virus globally. Noteworthy, Park et al. (2020) designed primers were implemented for traditional PCR and SYBR Green dye-based real-time RT-PCR, which methods required no probes. However, besides primer mismatch, sequence variations in probe recognition sites might also affect the efficiency of RT-PCR-based detection of COVID-19 providing false negative (unable to bind) or false positive (non-specific binding) results (Kamau et al., 2017; Nalla et al., 2020).

Analyzing only 180 SARS-CoV-2 sequences, Nalla et al. (2020) also reported primer-probe mismatches in strains of different countries with lower PCR sensitivity. But their investigation could not represent the global scenario due to limited sequences they considered. Strikingly, our study provides a global insight of N protein evolution and its consequences on primer-based detection of SARS-CoV-2 by RT-PCR method. Overall, global evolutionary insights on the primer-probe binding sites explore the undetected fragility of N gene-based nucleic acid detection of SARS-CoV-2, which warrant continuous monitoring and redesign of primer-probe sequences for efficient and accurate detection of COVID-19.

### 3.4 Amino acid deletions within the flexible SR-rich domain impact negatively c

Besides the synonymous and nonsynonymous mutations, our analysis explored 11 in-frame deletions of different ranged nucleotides across the N nucleocapsid sequences of SARS-CoV-2 originated from different countries worldwide (Table 5). The deleted nucleotide sequences distributed from 492 to 717 which fall into 3 major regions of N protein i.e. nt-positon ranges 492-494 (aa: 165), 578-633 (aa: 193-211) and 715-717 (aa: 239) (Table 5). Amino acid deletion at position 165 in the NTD RNA binding domain is not in the surface exposed region. Deletions of aa positioned 194-211 are in or close to the SR-rich linker region, and aa of position 239 is close to CTD domain and surface exposed (Fig 7). Surface exposed deletions near the CTD region may have a significant impact on nucleocapsid-RNA interaction and virus pathogenesis (Islam et al., 2020). Among the deletions, nucleotide deletion positioned 578-633 (aa: 193-211) found in multiple strains of multiple countries including Turkey, Bosnia, Australia, Malaysia, Germany, Scotland, England, Wales, and USA (Table 5, Supplementary Data 1). However, previously we explored 12 deletion-sites of ranged nucleotides in polyprotein (n = 9; NSP1:6, NSP2:1, NSP8:1, NSP15:1), ORF10 (n=1) and 3’-UTR (n=2) and 4 major deletion-sites in S protein while no deletions found among strains globally (Islam et al., 2020a; Rahman et al., 2020).

**Table 5:** Deleted positions in N protein of SARS-CoV-2.

| Nucleotide Position | Country | Accession number | aa position | Deleted aa |
| --- | --- | --- | --- | --- |
| 492-494 | Wales | EPI_ISL_460617 | 165 | T |
|  | Wales | EPI_ISL_468066 | 165 | T |
| 578-580 | England | EPI_ISL_463893 | 193 | S |
| 579-587 | Turkey | EPI_ISL_427137 | 194-196 | SRN |
|  | Turkey | EPI_ISL_490102 | 194-196 | SRN |
| 601-606 | Bosnia_and_Herzegovina | EPI_ISL_437226 | 201-202 | SS |
| 606-623 | Australia | EPI_ISL_478124 | 203-208 | RGTSPA |
|  | Malaysia | EPI_ISL_440977 | 203-208 | RGTSPA |
| 608-616 | Germany | EPI_ISL_425672 | 203-205 | RGT |
|  | Scotland | EPI_ISL_461586 | 203-205 | RGT |
| 620-631 | England | EPI_ISL_440146 | 208-211 | ARMA |
| 622-630 | Scotland | EPI_ISL_470172 | 208-210 | ARM |
| 623-625 | England | EPI_ISL_439913 | 209 | R |
| 625-627 | England | EPI_ISL_446738 | 209 | R |
|  | England | EPI_ISL_446708 | 209 | R |
|  | England | EPI_ISL_446700 | 209 | R |
| 626-634 | Wales | EPI_ISL_446655 | 208-210 | ARM |
|  | Wales | EPI_ISL_448809 | 208-210 | ARM |
|  | Wales | EPI_ISL_463145 | 208-210 | ARM |
|  | Wales | EPI_ISL_461507 | 208-210 | ARM |
|  | England | EPI_ISL_448135 | 208-210 | ARM |
|  | USA | EPI_ISL_434385 | 208-210 | ARM |
| 631-633 | England | EPI_ISL_460617 | 211 | A |
| 715-717 | Scotland | EPI_ISL_468066 | 239 | Q |
|  | Belgium | EPI_ISL_463893 | 239 | Q |

**Figure 7:**
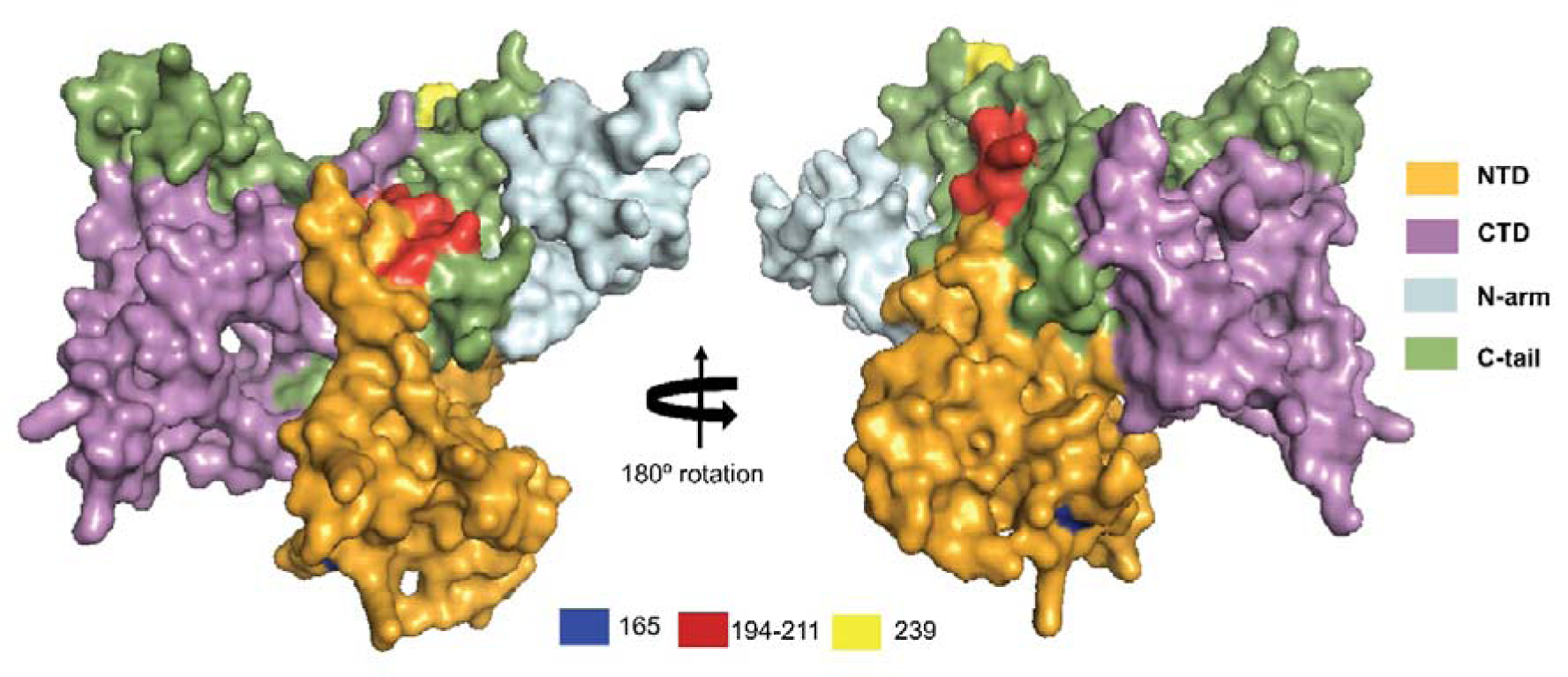
Deletion analyses of N protein of SARS-CoV-2. Deletion position of Nucleocapsid protein. 165 position is in the NTD RNA binding domain but not in the surface exposed. 194-211 are in the SR linker region and 239 is very near to CTD domain and surface exposed. Structures were visualized in PyMol v2.4.

These deletions may affect the virus-host interactions, immune-modulations, attenuation, and pathogenicity, by potentially influencing the tertiary structures and functions of the proteins (Islam et al., 2020b; Phan, 2020). Though, possible consequences of the deletions need to be observed to explore the immuno-pathogenic functions of the proteins and further implication in the development of effective therapeutics and prophylactics for SARS-CoV-2.

## 4. Conclusion

This study comprehensively explored the global evolution of SARS-CoV-2 nucleocapsid till 17 July, 2020. Our mutational analyses revealed 49.15 % nucleotide-level mutant strains, and 46.07 % of the mutant strains undergoing aa substitutions in their N protein sequences. We found mutational mismatches within primer binding sites particularly at 3’ end of the primer sets targeting N gene, which may result in inconsistent or higher false negative rates in detection tests of SARS-CoV-2. Furthermore, this study found significant impact of the amino acid group transversions along with deletion mutations on the tertiary structure and functions of the nucleocapsid of SARS-CoV-2. As N protein variation is high compared to the S protein of SARS-CoV-2, vaccine candidate targeting N will be challenging. Overall, this study warrants continuous monitoring of the ongoing evolution of the SARS-CoV-2 N protein for prophylactic and diagnostic interventions.

## Acknowledgments

The authors sincerely appreciate the researchers worldwide who had deposited and shared the complete genomes data of SARS-CoV-2 and other coronaviruses to GISAID (https://www.gisaid.org/).

## Data availability

This study utilized the SARS-CoV-2 genome sequences retrieving from the publicly available open database, GISAID. Detailed step by step methods are described in Mutation_analysis.pdf (https://github.com/SShaminur/Mutation-Analysis).

## Authors contributions

MSR, MRI, ASMRUA, II, MNH, and SA, conducted the overall study and draft the manuscripts. MSR finally compiled the manuscript. MMR, MS and MAH contributed intellectually to the interpretation and presentation of the results.

## Figures

**Supplementary Figure 1:**
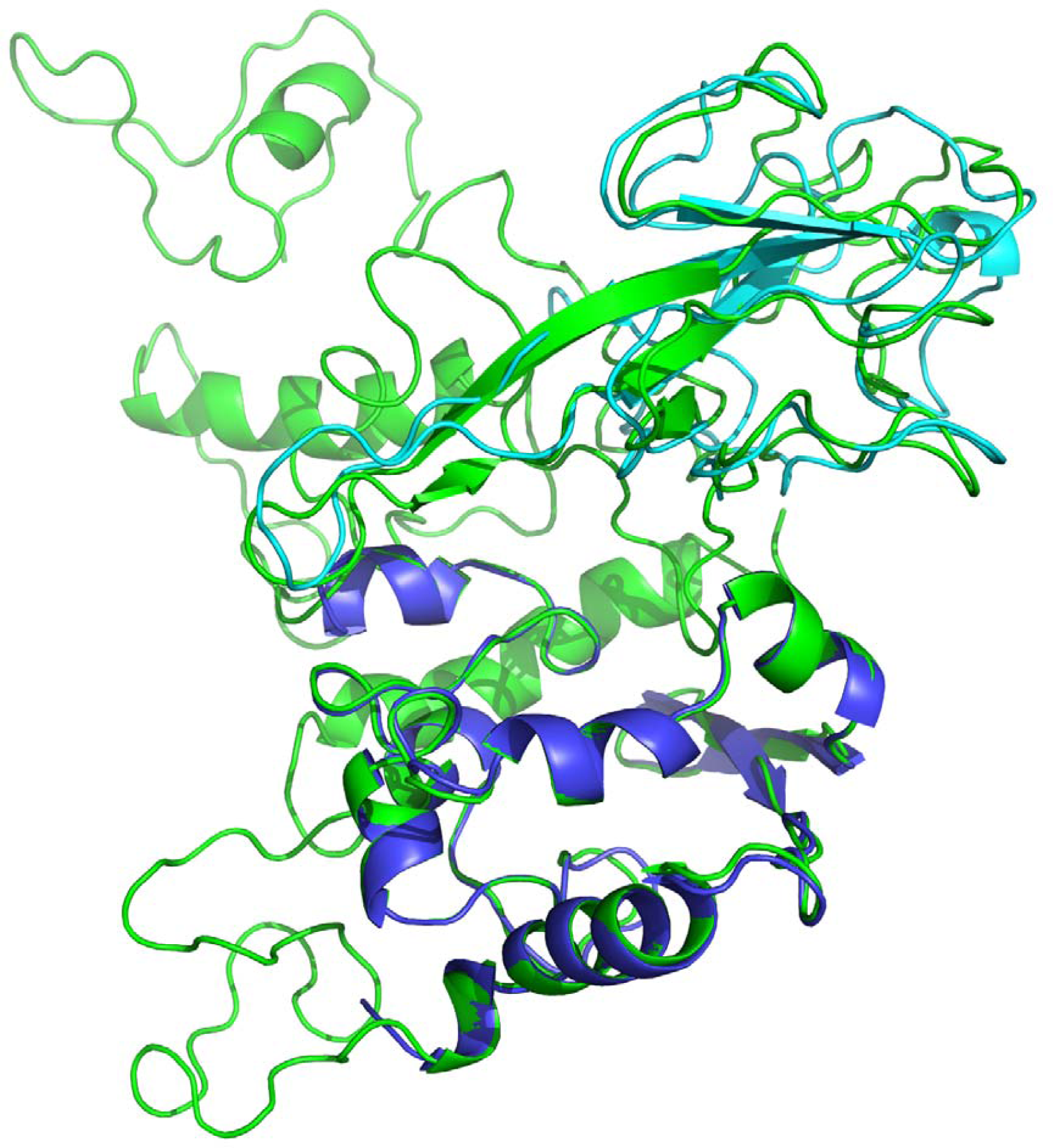
Structural alignment of the modeled 3D structure (green) with original NTD (cyan) and CTD (blue). Structures were visualized in PyMol v2.4.

